# Genetically distinct microenvironment determines cancer survival and response to therapy in mice

**DOI:** 10.64898/2026.07.10.737486

**Authors:** Mark A. Warner, Jennifer K. Sargent, Shawnna R. Farley, Beth L. Dumont, Muneer G. Hasham

**Author notes:** Corresponding author Muneer G. Hasham 600 Main Street, Bar Harbor, ME 04609.

## Abstract

Genetic uniqueness of the tumor microenvironment significantly influences cancer growth, survival, and response to therapy, independent of the cancer cell’s intrinsic properties or the adaptive immune system. Using genetically distinct *Rag1-/-* mouse models, this study shows that different strains exhibit varied tumor growth kinetics and survival outcomes when xenografted with identical leukemic and solid tumor cell lines. This study further highlights the critical role of the myeloid immune compartment and shows that disrupting both lymphoid and myeloid systems alters cancer progression. These results also reveal that the tumor microenvironment can permanently alter cancer cell phenotypes and significantly affect chemotherapy efficacy, as seen with Cisplatin’s varying effects across strains. These findings underscore the importance of considering genetic background in preclinical cancer models, suggesting that reliance upon a single mouse strain may lead to incomplete conclusions about cancer biology and treatment efficacy.

**SUMMARY STATEMENT:** Pre-clinical xenograft mammalian models are used to study human diseases. Here we report that the genetic uniqueness of the tumor microenvironment, independent of the immune system, can determine the fate of cancer progression, survival, and therapy response.

## INTRODUCTION

Cancer remains one of the leading causes of death worldwide, with its incidence rising in developing nations over recent decades (ARNOLD *et al*. 2017; SUNG *et al*. 2021; WILKINSON AND GATHANI 2022). Despite many therapy advancements, there is significant inter-individual variability in patient survival and disease progression in cancers resulting from the same genetic mutation (COATE *et al*. 2010; TALSETH-PALMER AND SCOTT 2011). This suggests that the microenvironment surrounding the cancer cell – and the host genetics underpinning its characteristics – may determine the outcome of the disease (ANDERSON AND SIMON 2020; DE VISSER AND JOYCE 2023). A microenvironment is comprised of the molecules, cells, and structures that support unique cells and tissues, such as the immune cells, stromal cells, signaling molecules, and vasculature surrounding a tumor (ANDERSON AND SIMON 2020). This definition can be broadened to consider the entire organism as the microenvironment for liquid cancers, such as leukemia. Abnormal cells, such as cancer cells, can alter their microenvironment which can in turn influence how cancer cells grow and spread (WU AND DAI 2017; DE VISSER AND JOYCE 2023). As such, the microenvironment poses an attractive target for improving our understanding how cancer cells form and developing new strategies for cancer treatment (WU AND DAI 2017; THAKKAR *et al*. 2020; XIAO AND YU 2021).

The relationship between cancer cells and their surrounding microenvironment is bidirectional: while cancer cells can remodel their environment, the microenvironment can also shape tumor growth and gene expression profiles (SARGENT *et al*. 2022; ELHANANI *et al*. 2023; HASHAM *et al*. 2024). In patients, unlike xenografted models, cancer cells, the microenvironment, and the immune system are all derived from identical genetic composition. Therefore, it is difficult to determine the exact contribution of the tumor microenvironment towards growth or response to therapy. In addition, a myriad of factors influences cancer growth and response to therapy, including the complexity of the human genome, adaptive immune response, living conditions, and lifestyle/environmental factors.

Mouse models provide an ideal solution to control not only an organism’s genetics, but also the environmental factors to which they are exposed, making them well-suited for translational cancer research (HOLLAND 2004; CHEON AND ORSULIC 2011; OLSON *et al*. 2018; BOSENBERG *et al*. 2023). Additionally, when performing genomic analysis on xenografted mouse models, one can computationally separate the human and mouse genomes or proteins, thereby making xenograft models an excellent vehicle to probe the genetics of the host versus the cancer (JUNG *et al*. 2018; HASHAM *et al*. 2024). These models allow for controlled manipulation of tumor–host interactions, enabling researchers to study how the tumor microenvironment, tissue context, and immune components collectively influence treatment response and disease outcomes. However, to truly understand the extent genetics can govern the microenvironment, syngeneic and graft model systems must be deconstructed and tested strictly on the contribution of the tumor-environment, independent of the immune system and with the same cancer line.

Therefore, we developed and tested multiple genetically diverse *Rag1* knockout mice (*Rag1-/-*), which inhibits the development of the adaptive immune system and allows human cancers to be xenografted. *Rag1-/-* arrests the development of T and B lymphocytes at an early stage, preventing maturation in, and exit from, the bone marrow (FUGMANN 2001). As previously published, when the same clonal human solid tumor cell line (MDA-MB-231, derived from a triple negative breast cancer patient) was xenografted subcutaneously in these unique strains, each strain exhibited different growth rates (SARGENT *et al*. 2022; HASHAM *et al*. 2024). The same was observed with a chronic lymphocytic leukemia human cell line (MEC1). and a mouse glioblastoma cell line (GL261) when engrafted into *Rag1-/-* mice, indicating that genetically different tumor microenvironments influenced the growth kinetics of the cancer cells independent of both the adaptive immune system and the intrinsic genetic factors of the cell line used (SARGENT *et al*. 2022; HASHAM *et al*. 2024). Proteomic profiling of the xenografted tumor cells revealed unique protein expression signatures suggestive of cancer cell evolution to more aptly suit to the tumor microenvironment (HASHAM *et al*. 2024).

Our recent studies demonstrate that the genetic makeup of the microenvironment can significantly influence tumor behavior, independent of adaptive immunity (SARGENT *et al*. 2022; HASHAM *et al*. 2024). Using *Rag1-/-* mice, which lack T and B lymphocytes, we found that the same cancer cell line exhibited strikingly different growth kinetics and protein expression profiles when xenografted into genetically distinct host strains (SARGENT *et al*. 2022; HASHAM *et al*. 2024). In fact, the growth of three different cancer cell lines across multiple *Rag1*-deficient strains revealed a broader range of tumor kinetics driven by host background than by tumor type itself, underscoring the dominant role of microenvironmental genetics in modulating tumor progression and disease outcomes (SARGENT *et al*. 2022; HASHAM *et al*. 2024).

In this study, we further interrogate the overarching hypothesis that the tumor microenvironment plays a more significant role in cancer progression than the cancerous cells themselves, and that this microenvironment is directly governed by the genetics of the host. Specifically, we seek to determine **(a)** whether the tumor microenvironment influences survivorship in different strains of mice with the same xenografted cancers, **(b)** if cancer growth and survival is influenced by the myeloid compartment of the tumor microenvironment, **(c)** how the genetically distinct tumor microenvironment changes the phenotype of the cancer cell, and **(d)** whether genetically different microenvironments influence response to therapy independent of the adaptive immune system. Together, these investigations aim to define how genetic variation in the tumor microenvironment shapes cancer progression and treatment response, independent of both the cancer cell itself and the adaptive immune system.

## RESULTS

### Survival of genetically distinct immunodeficient (*Rag1-/-*) mouse strains with leukemia

We previously demonstrated that identical human or mouse cancer cell lines, grown in culture and then xenografted into five genetically distinct mouse strains, exhibit markedly different growth kinetics depending on the host background (SARGENT *et al*. 2022; HASHAM *et al*. 2024). Furthermore, these mice were immunodeficient in both T- and B- lymphocytes due to their *Rag1* nullizygosity (SARGENT *et al*. 2022; HASHAM *et al*. 2024). This data indicates that the genetic microenvironment in which the cancer is engrafted, independent of the adaptive immune system and the intrinsic nature of the cancer cell, determines the growth of both solid and liquid cancers. However, it is important to note that, due to ethical reasons, xenografted solid tumors can only be allowed to grow to a certain size (e.g., 1000 mm^3^) before the mouse must be euthanized. *Therefore, it is important to recognize that the size of the tumor does not necessarily indicate the inability of a certain strain of mouse to sustain the cancer without symptoms*. In general, a patient does not typically die from the size of a tumor, but rather what the tumor does to the rest of the body, and thus tumor size alone is not fully reflective of the disease. However, a leukemia model can be used to address the question of whether the tumor microenvironment also influences survivability apart from tumor growth.

To this end, three different strains of *Rag1*-/- mice were xenografted with MEC1, a chronic lymphocytic leukemia cell line, to generate previously validated xenograft models. The MEC1 cell line carried a luciferase cassette which enabled the longitudinal assessment of the tumor burden in the same mouse by live imaging. Since a leukemia xenograft model is not burdened by the physical size of a tumor such as in a subcutaneous xenograft, this leukemia xenograft model is an ideal system to test survival in genetically distinct strains. Unlike solid tumors, leukemia engages the entire organism as its microenvironment, making it particularly well-suited to interrogate how host genetics shape cancer progression (HOPKEN AND REHM 2019; CIANTRA *et al*. 2025)

To capture an expansive allelic divergence in our studies, six of the parental strains of the Collaborative Cross/ Diversity Outbred platforms (129S1/SvImJ (129R), A/J (A/JR), C57BL/6J (B6R), CAST/EiJ (CASTR), NOD/ShiLtJ (NR), and PWK/PhJ (PWKR)) with *Rag1-/-* were used. The Collaborative Cross/ Diversity Outbred together capture, on average, 90% of the known allelic divergences across the mouse genome and approximately 0.25 million years of evolutionary divergence (**Figure 1A**) (ROBERTS *et al*. 2007; PHIFER-RIXEY *et al*. 2020). These mice also capture the most commonly used subspecies of *Mus musculus*—the *domesticus, castneus, and musculus* (**Figure 1A**). In addition, our previous studies showed that BALB/cJ *Rag1-/-* (BAR) mice had a tumor growth that was in between the fastest and slowest growing strains, thus was added to the panel. NSG^TM^ and J:NU (athymic nude mice) were added as control strains, as they are commonly used in xenograft growth, survival, and efficacy studies. Their inclusion provides well-characterized reference points for comparison against the genetically diverse strains in our panel (HOFFMAN 2017; UTHAMANTHIL *et al*. 2017; JUNG *et al*. 2018; BOSENBERG *et al*. 2023).

**Fig. 1.**
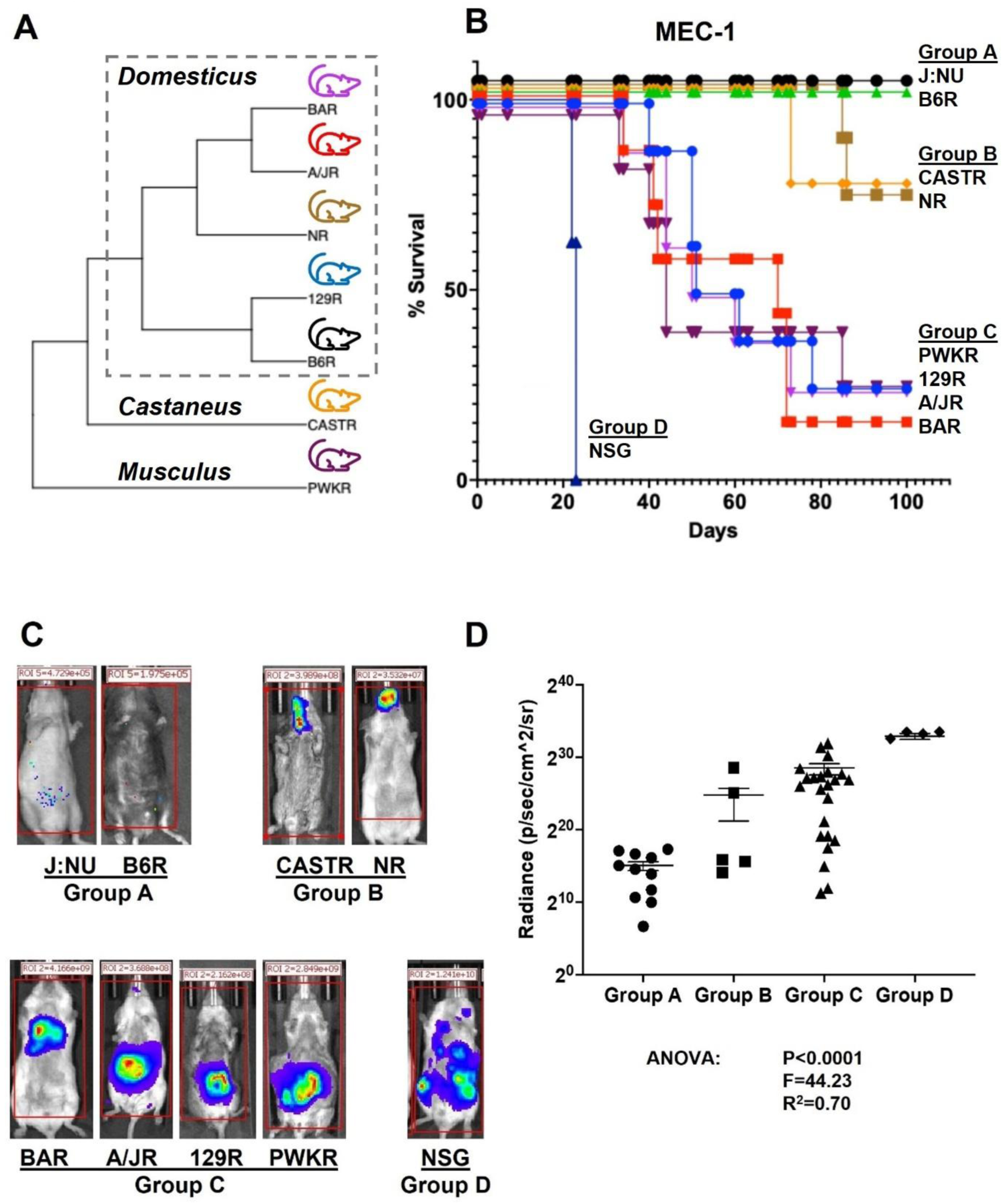
Genetically distinct micro-environment determines the growth kinetics of leukemia. **(A)** A cladogram showing the evolutionary divergence of the strains used in the study. Strains from top to bottom: BALB/cJ *Rag1-/-* (BAR), A/J *Rag1-/-* (A/JR), NOD/ShiLtJ *Rag1-/-* (NR), 129S1/SvImJ *Rag1-/-* (129R), C57BL6/J *Rag1-/-* (B6R), CAST/EiJ *Rag1-/-* (CASTR), and PWK/PhJ *Rag1-/-* (PWKR). **(B)** Survivorship by strain. MEC1 cells with a luciferase cassette were injected via intracardiac injections in 9 different *Rag1-/-*mouse strains to induce leukemia. The survival curve is presented as percentage of survival due to different sample sizes. The endpoint for euthanasia was determined by a <2.5 Body Condition Score (BCS), dehydration, lethargy, or animal care staff reporting one of these conditions. After 100 days, the remaining mice were imaged before euthanasia. All strains consisted of 4 female and 4 male mice except for A/JR and NR females, and BAR and PWKR males which all had 3 mice. **(C)** An image of each strain prior to euthanasia is shown as a representative. Group A shows very little to no measurable signals. Group B shows signals localized to the sinuses and lymph nodes around the neck. Group C shows high signals localized in the body cavity and organs. Group D shows the highest signals, dispersed throughout the body cavity. **(D)** A scatter plot with Radiance (p/sec/cm²/sr) is presented in the previously established groups. ANOVA: P<0.0001, F=44.23, and R²= 0.70.

To test whether the genetic diversity of the tumor microenvironment influences survival outcomes, we xenografted mice via intracardiac injection of 2 x10^6^ MEC1 cells per mouse. Equal numbers of male and female mice were included, as prior studies showed no significant sex-based differences in leukemia growth (SARGENT *et al*. 2022). Although tail vein injections are a standard method for leukemia xenografts, strain-dependent differences in vein visibility and accessibility would introduce unwanted variability. Intracardiac injection was therefore selected to ensure consistency across all mouse strains.

Survival analysis of the genetically distinct mouse panel revealed four distinct patterns of disease progression (**Figure 1B**). *Group A* consisted of J:NU and B6R mice, all of which survived the full 100-day study. *Group B* contained CASTR and NR mice that showed a majority in latency of 73 or more days before the first mice fell ill. Illness was defined as the lack of ambulation, the inability to groom or feed, or definitive signs of general malaise/morbidity. *Group C* consisted of PWKR, 129R, A/JR, and BAR strains, which had a staggered latency starting at 33 days with only 1 or 2 animals surviving to the end of the study. *Group D* contained NSG control mice, which all fell ill early in the study (at 22-23 days), although there is a formal possibility that this was due to factors other than the leukemia. Since the MEC1 cells carried a luciferase cassette, we measured the amount of leukemia in the mice via live *in vivo* imaging (**Figure 1C and D**). The overall leukemia burden resulting from MEC1 cell administration was comparable across groups; however, leukemia progression differed significantly between strains, as tested by Analysis of Variance (ANOVA), indicating that differences in survival were attributable to host-dependent responses to disease rather than variation in initial tumor load or other causes.

These results strongly suggest that the cancer microenvironment determines leukemia survivorship. Surprisingly, some mouse strains that are genetically distant based on single nucleotide polymorphisms (SNP, **Figure 1A**) profiles nonetheless displayed similar leukemia survival outcomes. For instance, NR and CASTR both exhibit moderate survivorship (Group B), while 129R and PWKR showed significantly lower survivorship in Group C, despite SNP-based analyses predicting greater similarity between 129R and NR. These findings suggest that leukemia-related survival phenotypes may be shaped by genetic factors beyond those captured by SNP variation alone, emphasizing the importance of utilizing genetically different xenograft model systems.

### Contribution of the genetically distinct myeloid compartment in cancer growth

The immune system is comprised of the lymphoid and the myeloid compartments (PARHAM AND JANEWAY 2015). While it is well established that xenografts can grow in the absence of the lymphoid compartment, the intact myeloid system can still prevent tumor growth (SARGENT *et al*. 2022). Previously and in **Figure 1**, we tested the ability of genetically different lymphodeficient (*Rag1-/-*) mice to sustain leukemia. However, since the myeloid component can influence the growth of cancer and ultimately survival in a leukemia model, both the lymphoid and the myeloid components need to be disrupted. A viable way to induce such immunodeficiency is by removing interleukin receptor 2 gamma subunit (*Il2rg-/-*), as it has been shown in xenograft models (CAO *et al*. 1995). Since Group A has low engraftment, and Group D is a control group for the survey experiment, they were not used. Therefore, we chose to proceed in follow up studies using both strains from Group B (CASTR, NR) and a representative of Group C (PWKR) to determine whether the myeloid component contributes to leukemia growth, and ultimately, survival. In addition, the choice of strains was based on the maximum divergence in the panel (**Figure 1A**). Therefore, we obtained NRG, CRG and PRG mice, where the first letter denotes the strain (**N**OD/ShiltJ, **C**AST/EiJ, or **P**WK/PhJ), the second letter denotes ***R****ag1-/-* and the third letter denotes *Il2r**g**-/-*, to assess the impact defective lymphoid and myeloid compartments on cancer outcomes. MEC1, CCRF-SB (an acute lymphoblastic leukemia cell line), and the solid tumor line MDA-MB-231 (a triple negative breast cancer cell line) were xenografted in NRG, CRG, and PRG strains to test the extent that the myeloid immune system influences leukemia growth, mouse survival, and solid tumor growth, furthering the scope of our cancer panel and improving translational potential of these findings.

The disruption of the myeloid component, in addition to lack of the lymphoid compartment, reversed the survival phenotype of PRG and NRG mice with MEC1 xenografts. While NR showed higher survival than PWKR (**Figure 1B**), PRG could sustain the leukemia significantly better than NRG mice (**Figure 2A**). The CRG mice did not show engraftment with MEC1 cells, although it is unclear if this was due to an inability of the cells to grow in the CRG microenvironment or an issue with the xenograft itself. When comparing the spleen mass between NRG and PRG mice, splenomegaly was significantly higher in PRG mice, suggesting a higher tumor load since MEC1 cells reside mostly in the spleen (**Figure 2B**) (WILSON *et al*. 2019). MEC1 in the spleen or bone marrow was measured as the percentage of human CD19 using flow cytometry marker for this xenograft, which was not different between the two strains (**Supplementary Figure 1**). The larger spleens in PRG mice reflects a higher total amount of human CD19 in PRG spleens than in NRG spleens. When another leukemia cell line CCRF-SB (from American Tissue Culture Collection, ATCC Cat no. CCL-120) was tested, a similar result was observed (**Figure 2C and D**). CRG survival was comparable to NRG, which mirrored the results when the myeloid compartment was intact (**Figure 1**, both are Group B). This data showed, at least for the PWK/PhJ strain, that the disruption of the myeloid component enhanced its leukemia burden carrying capacity.

**Fig. 2.**
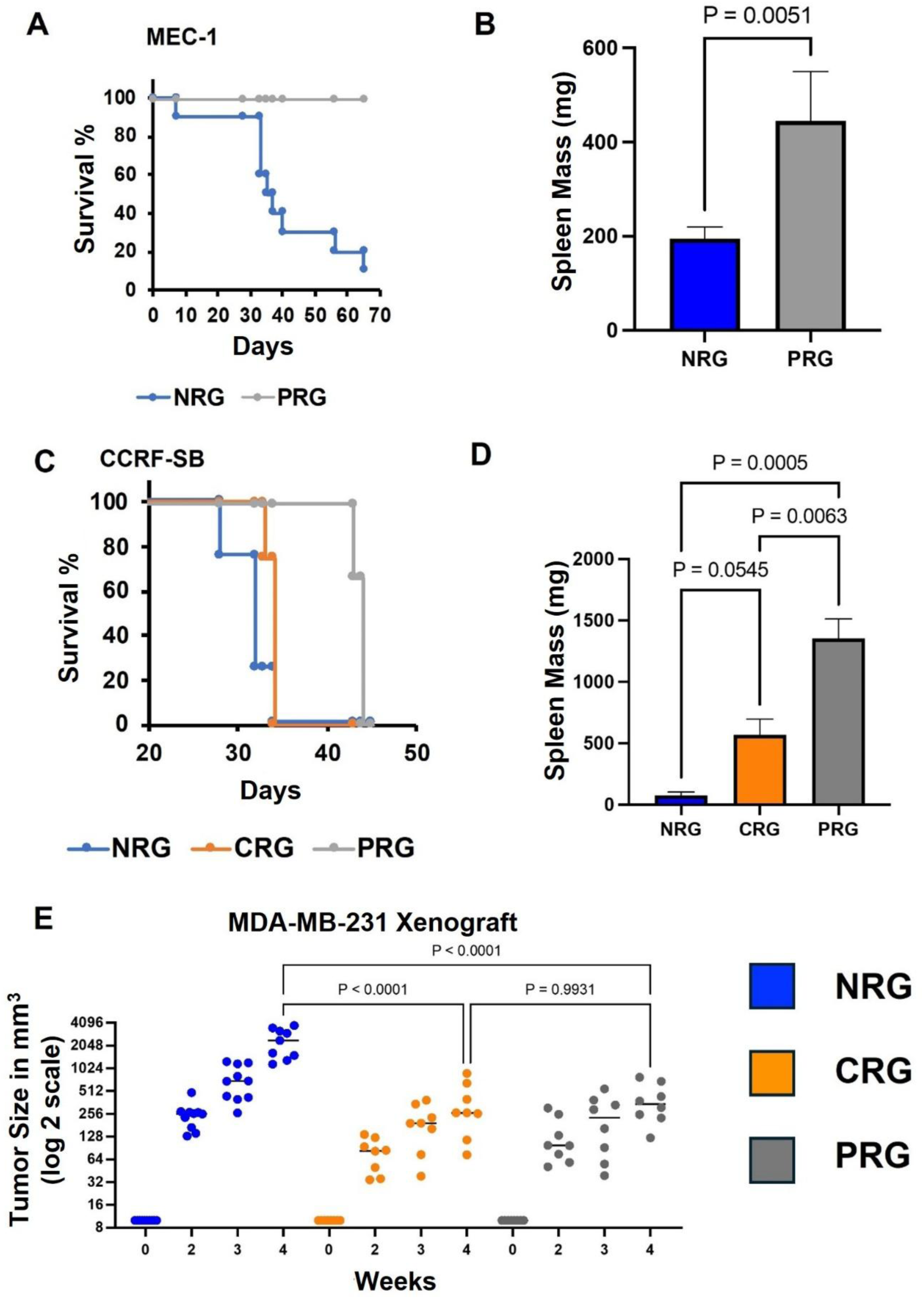
The myeloid component of the micro-environment influences survival.(A) Using the same endpoint criteria as in Figure 1B, a survival curve is shown with NRG and PRG mice, which were injected with MEC1 Luc via tail vein injection into 5 female and 5 male NSG mice, and 1 female and 2 male PRG mice. NRG mice had a latency of 28 days, with most requiring euthanasia by 40 days and only one surviving beyond 56 days. All the PRG mice required euthanasia between 65 and 69 days. **(B)** Average spleen mass is shown in milligrams **(C)** This survival curve shows CRG, NRG, and PRG mice injected via tail vein with CCRS-FB. 2 females and 2 males of each strain were used except for PRG, which only shows 1 female. Here we see a similar trend, with NRG mice requiring euthanasia earlier and PRG mice surviving longer, but with limited intra-group variability compared to NRG. CRG mice showed an intermediate phenotype, with a later latency of disease onset for the first mouse than in NRG mice, limited variability like PRG mice, and the last survival animal requiring euthanasia at the same time as the last NRG mouse. **(D)** Average spleen mass is shown for the CRG, NRG, and PRG mice that were engrafted with CCRF-SB from Figure 2C. NRG mice still exhibited small spleens averaging 79mg, CRG had enlarged spleens averaging 570mg, and PRG had extremely enlarged spleens averaging 1352mg. **(E)** In addition to liquid models of leukemia, we tested the growth of a solid tumor, a triple negative breast cancer, MDA-MB-231 Luc. This scatter plot shows the growth over the course of 4 weeks in log2 scale of CRG, NRG, and PRG mice which were xenografted subcutaneously. The 4 week endpoint was determined by NRG tumors exceeding 1000mm^3^.

We next asked whether myeloid disruption also affects the progression of solid tumors. Therefore, NRG, PRG, and CRG mice were xenografted subcutaneously with MDA-MB-231, a solid tumor cell line, that had previously been used to test the effects of the genetic distinct tumor microenvironment (SARGENT *et al*. 2022). NRG mice experienced significantly greater tumor growth compared to PRG and CRG strains, suggesting that while the PRG microenvironment may favor leukemia, it is less permissive to the growth of this solid tumor (**Figure 2E**).

### Pro-inflammatory cytokine IL-1B from the microenvironment and not the immune system determines leukemic expansion

Inflammation is a major source of cancer growth (HANAHAN 2026). However, the source of pro-inflammatory cytokines such as IL-1B cannot be easily studied since both the immune system and the micro-environment could potentially contribute to the secretion of these molecules. In our previous studies with *Rag1-/-* mice, we observed that the strains that had the best growth has the largest change in pro-inflammatory cytokines such as IL-1B (SARGENT *et al*. 2022). These results suggested that the pro-inflammatory cytokines controlled the growth of the tumor. We hypothesized that reduction of pro-inflammatory cytokine such as IL-1B would reduce the growth. Therefore, we used PRG mice as a model to test this hypothesis (**Figure 3**). We xenografted mice with CCRF-SB cells similar to **Figure 2C and 2D** and then 10 days after we intraperitoneally injected 10µg of IL-1 receptor alpha chain inhibitor (IL-1Ra InVivoKine, AdipoGen, Cat # AG-40B-0254-C050) in 100µL of sterile saline twice a week for two weeks. This inhibitor should block IL-1B binding to the cells that promote cancer growth. Using In vivo imaging, it was observed that leukemia was significantly reduced in the presence of this inhibitor, suggesting that pro-inflammatory cytokines, such as IL-1B, from the non-immune component of the micro-environment controls cancer progression. This was confirmed by spleen mass and flow cytometry of the blood by staining for leukemia cells (PE Mouse Anti-Human CD45 BD Pharmingen Cat # 561866) (**Figure 3B and 3C**). There was a reduction of leukemic cells in the bone marrow and spleen, albeit not significantly, suggesting that these niches might require further treatment to obtain complete ablation.

**Fig. 3.**
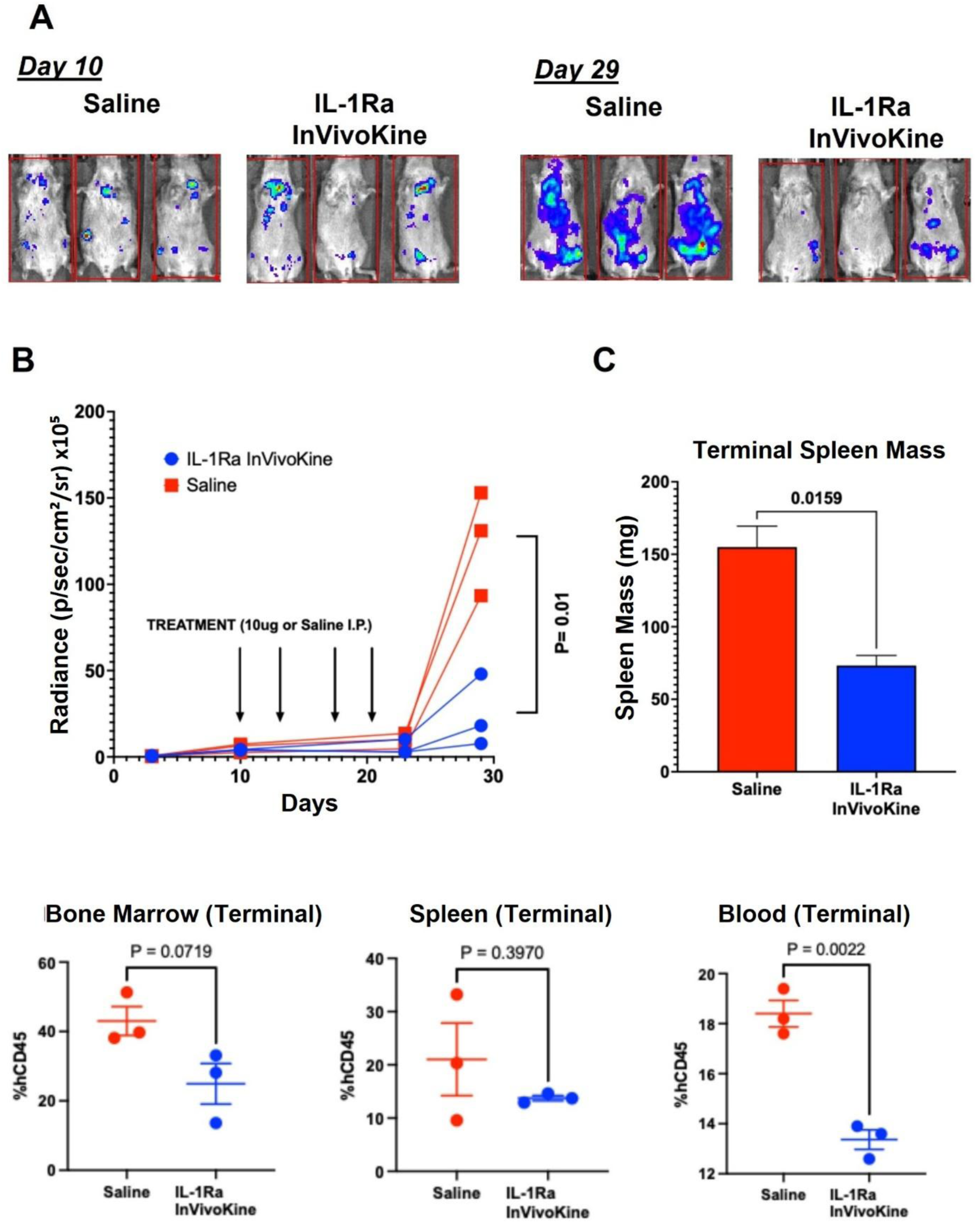
Pro-inflammatory cytokine IL-1B from the microenvironment determines leukemic expansion. **(A)** CCRF-SB Luc was given to 6 female PRG mice via intracardiac injection and imaged after 10 days to test growth. Both cohorts showed comparable signal, so treatment of IL-1Ra InVivoKine, an IL-1B inhibitor, was administered via IP injection twice a week for 2 weeks and were imaged at 23 days post intracardiac injection, and again at 29 days post intracardiac injection. Day 10 images are on the color scale Min = 7.50e3 and Max = 5.62e4, and Day 29 are on the color scale Min = 4.25e6 and Max = 6.64e7; blue is lowest and red as highest signal. The graph shows the radiance for imaging points and treatment days. **(B)** Terminal spleen mass is shown in mg, IL-1Ra InVivoKine treated spleens show less splenomegaly than the saline control treatment group. **(C)** Flow cytometry shows human CD45 stained cells from the spleen, bone marrow, and blood. Only the blood shows a statistically significant difference, P = 0.0022 aligning with the imaging.

### The genetics of the tumor microenvironment alter cancer cell phenotypes

While we have shown both previously and in this study that the growth kinetics of a given cancer cell line are dictated by surrounding the microenvironment (**Figures 1 and 2**), it remains unclear whether these effects are transient or permanent: *Does the cancer cell adapt its phenotype in response to the microenvironment? Does it undergo stable, heritable changes? Or does the microenvironment selectively support a pre-existing subclone within the population?* This <u>adaptation</u> versus <u>evolution</u> question is important to determine and to truly understand the role of the tumor microenvironment in the growth of cancer. To answer this question, we turned to solid tumor models, as leukemia cells are distributed throughout the bloodstream and lymphoid organs, making it difficult to isolate a specific site for interrogation without introducing bias in the observed phenotype. For instance, cancer cells in the bone marrow may be readily adapting to the microenvironment, while the cells in the spleen have been selected for a certain phenotype. While this would be interesting, it would require investigation beyond the scope of this study. To move forward, we focused on solid tumor models with the most extreme phenotypes. Based on prior studies, B6R was selected as the model with the slowest tumor development, and NR as the model with the fastest tumor growth (SARGENT *et al*. 2022).

To test whether cancer cells are merely adapting to the microenvironment or undergoing permanent changes or clonal selection, we first xenografted MDA-MB-231 clonal cancer cells in both NR and B6R mice. Our previous work has demonstrated that the NR microenvironment is growth inducive and B6R is growth inhibitive, the latter primarily due to massive collagen deposits within the tumor (SARGENT *et al*. 2022). When the tumors grew, we passaged them in equal amounts into both NR and B6R naïve mice (**Figure 4A**). Tumor size was measured using calipers over a period of 32 days. To assess whether cancer cells adapt to their microenvironment or are shaped by it, we first passaged the MDA-MB-231 cell line from a growth inducive environment (NR) and into a growth inhibitive environment (B6R). Tumors derived from this passage exhibited no difference in growth kinetics of the tumors compared to those maintained in the growth-permissive NR environment (**Figure 4B**). Similarly, cells passaged from a growth inhibitive environment (B6R) into a growth inducive one (NR), showed only a minor, non-significant difference in tumor size and growth compared to those continuously passaged in B6R (**Figure 4B**). Notably, tumors that originated in NR grew significantly larger than those that originated in B6R, regardless of the environment into which they were subsequently passaged (P≤0.0004, **Figure 4B**). These findings suggest that xenografted cancer cells do not adapt dynamically to microenvironments, rather the host microenvironment is the predominant driver of the effect, either selecting for or supporting tumor growth depending on its intrinsic permissiveness.

**Fig. 4.**
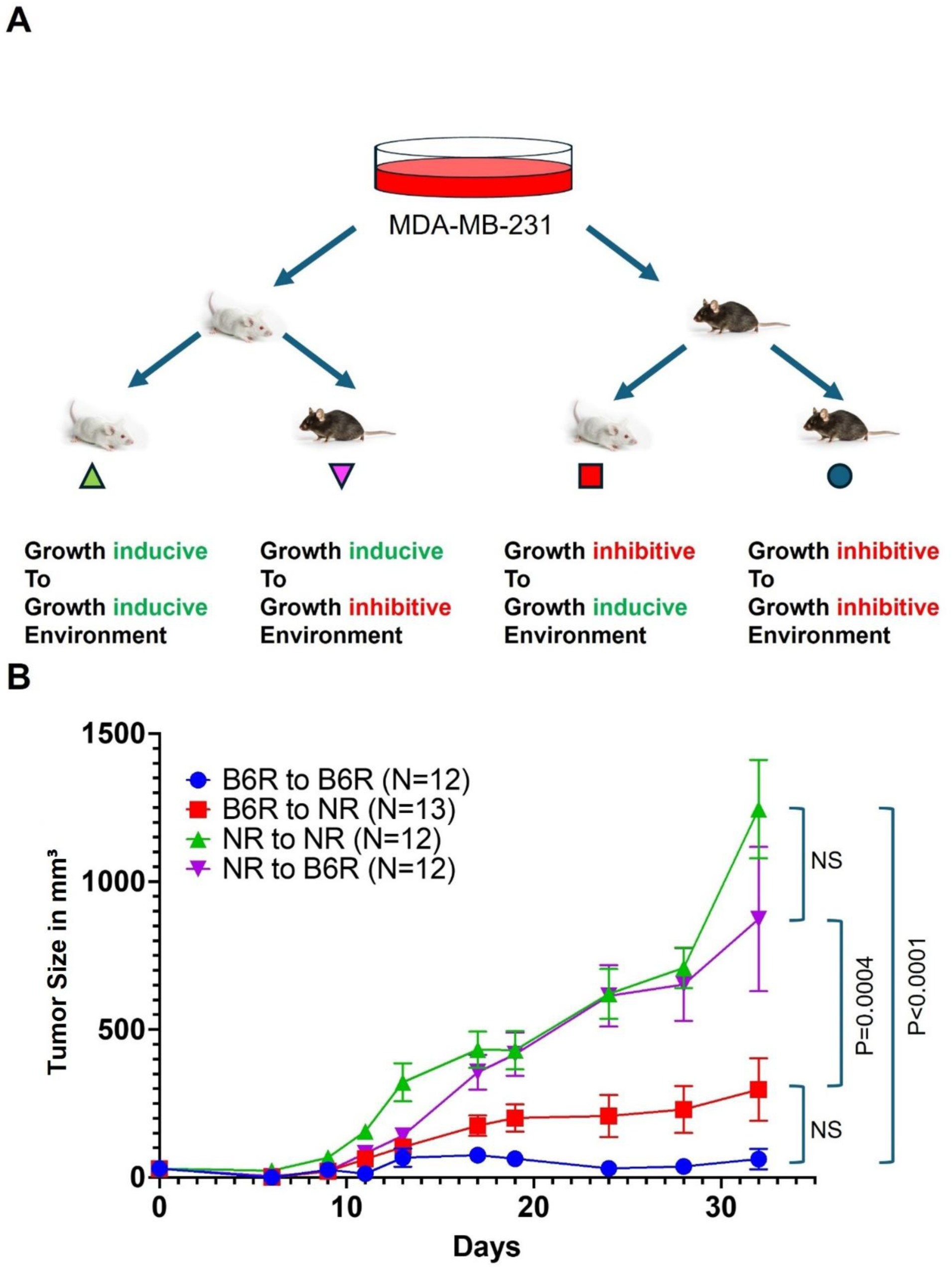
Evolution of a xenografted cancer is influenced by the genetic micro-environment. **(A)** To test if the initial host environment affects tumor growth in a future passage, we grew the same MDA-MB-231 Luc cell line in NR and B6R mice. Once the tumor reached 350mm^3^ in a strain, it was passaged into 12 mice from each strain. **(B)** The histogram of the average growth of the passaged tumors over the course of 32 days measured in mm^3^.

### Genetics of the microenvironment drive chemotherapy treatment efficacy

While understanding how genetically distinct tumor microenvironments influence cancer growth kinetics is biologically important, the most clinically critical phenotype remains the response to therapy. It is well established that the same treatment can yield vastly different outcomes across different patients. However, because a patient’s tumor shares the same genetic origin as the host, disentangling the specific contribution of the tumor microenvironment to therapeutic response remains a significant challenge. Additionally, tumors are often excised after resistance has already emerged, introducing further complexity due to sub-clonal heterogeneity. To disentangle these factors, xenograft mouse models afford us the ability to separate/differentiate the genetic contributions on the microenvironment to tumor growth.

For this experiment, we again leveraged the solid tumor system, since models such as B6R showed low xenograft growth in leukemia models (**Figure 1**), making it difficult to detect subtle differences between treated and control groups. We selected the slowest and fastest growing strains (B6R and NR, respectively), as previously determined (SARGENT *et al*. 2022). We also tested 129R, another strain shown to support rapid tumor growth in earlier studies (SARGENT *et al*. 2022; HASHAM *et al*. 2024). All three strains were xenografted with MDA-MB-231, which we and others have demonstrated to reliably grow in these genetic backgrounds. When the tumors reached at least ∼30 mm³, the mice were treated with Cisplatin, a common chemotherapeutic agent, three times a week for 21 days at 2 mg/kg I.P (PERSE 2021). Cisplatin is a DNA cross-linker and interferes with cell proliferation, and it is well documented to be efficacious in suppressing tumor growth in many cell lines, including MDA-MB-231 (PRABHAKARAN *et al*. 2013). The tumors in vehicle treated NR mice grew up to ∼20 fold but with Cisplatin the tumors grew only ∼3 fold. In B6R, vehicle treated tumors grew on average ∼3 fold and the Cisplatin treated tumors ∼2 fold. Treatment with Cisplatin showed an opposing effect within the tumors of 129R mice: after 21 days, tumors in vehicle treated mice grew ∼3.3 fold, and Cisplatin treatment resulted in > ∼6-fold increase in tumor growth (p=0.04) (**Figure 5**). Together, these data suggest that the genetic makeup of the tumor microenvironment, which includes the ability of tissues of different strains to metabolize Cisplatin differently, not only governs cancer growth but also plays a decisive role in shaping therapeutic response, even when the tumor itself remains genetically identical.

**Fig. 5.**
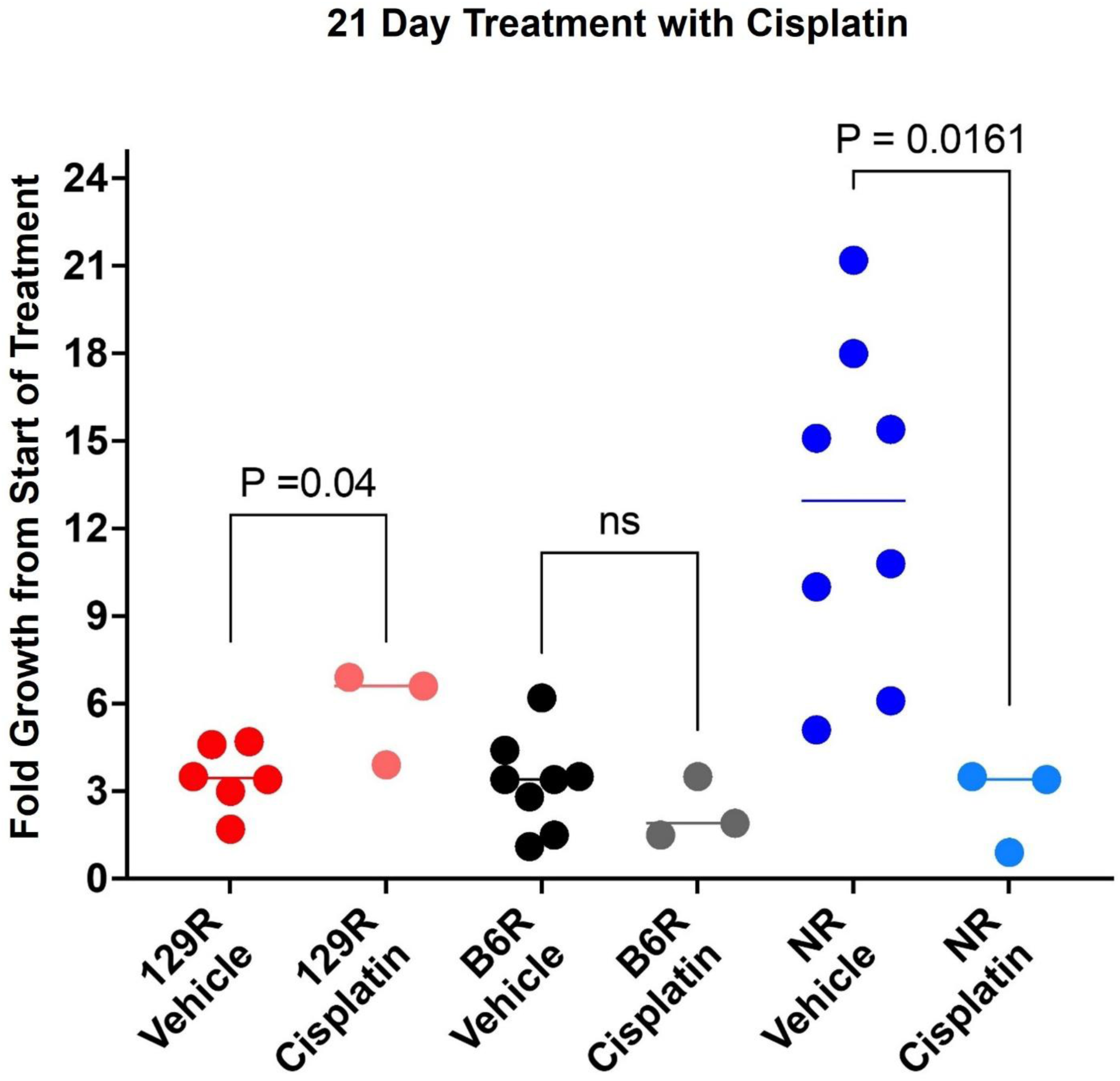
Therapy is affected by the genetic micro-environment. The effect of Cisplatin in genetically distinct mice engrafted with MDA-MB-231 Luc. Here, we show the fold growth of MDA-MB-231 Luc cells between 1 week post injection prior to Cisplatin treatment and 21 days post treatment with Cisplatin vs saline injected IP at 2mg/kg, 2 times per week. Mouse weight in grams x 2 was the volume injected IP for proper dosage in µL of Cisplatin in suspension of saline.

## DISCUSSION

In this study, we used both solid and liquid cancer models to understand how the tumor microenvironment influences cancer cell growth, therapeutic response, and overall survival across genetically distinct mouse strains. Numerous studies have looked into the role of tumor microenvironment in both biological models and patient samples (ANDERSON AND SIMON 2020). However, directly studying the tumor microenvironment in patient samples proves challenging. Patient-derived samples are inherently complex due to high genetic variability and the influence of the patient’s immune system, which cannot be controlled to the same extent as in genetically modified animal models. Because both the tumor and its surrounding microenvironment share the same genetic origin in patients, it becomes difficult to disentangle the specific contribution of the microenvironment to cancer progression or treatment outcome.

Previously and herein, we have developed and validated a mouse model system that can more robustly decipher the role of the tumor microenvironment. However, there are limitations with this model system: for instance, survival can only be studied with liquid tumor xenografts, as solid tumors cannot be allowed to grow beyond a certain size for ethical reasons. In order to produce metastatic lesions, many solid tumor models need to reach beyond the standard ethical size threshold, limiting the utility of this model. Although tumor extractions of these metastatic models can be performed, inherent variability in this *in vivo* system and an inability to control for significant mechanical and handling stressors may compromise the integrity of the animal and the metastatic tumor growth being evaluated. Importantly, it is also imperative to note that organismal survival rates and cancer growth kinetics are not equivalent parameters. It has been observed, commonly in chronic lymphocytic leukemia and benign hyperplastic growth patients, that the tumor burden is not correlated with their survival, alluding to the critical role of the microenvironment in governing patient survival (PEKTAS *et al*. 2025). Our previous work demonstrated the clear role of genetic background plays in cancer growth rates (SARGENT *et al*. 2022), and this study further characterizes the explicit role of genetic background in determining survival outcomes in xenograft cancer models.

In this study, we assessed the impact of growth permissive and growth inhibitive microenvironments in one solid tumor and two liquid tumor models. In addition to proteomic profiling of these microenvironment conditions, we also tested the efficacy of Cisplatin in a variety of *Rag 1-/-* strains, demonstrating a clear impact of both microenvironment and genetic context on treatment efficacy. It is important to note that the cell lines we have used are well-established in the field and adequately represent the biological processes associated with these cancers in patients, thus enabling the revelation of biological patterns that emerge due to genetic distinctness of the host. Despite the utility of these model systems, caution needs to be observed when generalizing the patterns observed in specific strains and cancer cell lines as directly representative of the biology and treatment responses for all cancers. Future studies will aim to test different classes/types of the same cancer, and whether the microenvironment plays a critical role in determining outcome or therapy response in those models as well. For example, while here we have demonstrated the impact of genetic background and microenvironment in the triple negative breast cancer cell line MDA-MB-231, future studies would determine whether a similar pattern emerges with a hormone dependent MCF cell line or comparable patient derived xenograft at various stages of cancer progression.

An intriguing feature was observed when the myeloid component of the immune system was disrupted in the leukemia studies. PWKR mice exhibited shorter survival than NR and CASTR when xenografted with MEC1 cells. However, when *Il2rg* was genetically removed from these strains, PRG mice showed longer survival and could carry a higher tumor burden than NRG or CRG. *Il2rg* is an essential component of IL-2 and other cytokine receptors, and its disruption targets the myeloid component of the immune system, particularly NK cells (CAO *et al*. 1995). Thus, our results have revealed that the genetic differences of the myeloid component of the immune system may play a crucial role in xenograft rejection, completely independent of the adaptive immune system. Therefore, these features make PRG an ideal mouse model for studying leukemias, especially in efficacy testing of slow-acting compounds. Unlike NRG or other commonly used strains, the extended disease latency allows sufficient time for delayed therapeutic effects to emerge.

Apart from studying cancer growth and effects, xenografted mouse models are highly pragmatic in pre-clinical testing, particularly in their ability to validate new compounds, molecules, or combination therapies for potential efficacy in human patients (BOSENBERG *et al*. 2023). Yet, there is an overarching tendency in the field to only evaluate efficacy using one or two strains of mice prior to determining if a particular compound or therapy is rejected or accepted as a viable for further testing. We have previously demonstrated that patient derived xenografts may fail in commonly used mouse strains such as NSG, but may xenograft very well in other *Rag1-/-* strains (HASHAM *et al*. 2024). Here, we show highly variable treatment outcomes, as Cisplatin treatment exhibited three distinct effects in three different strains. Cisplatin is a platinum compound that binds to DNA and upon replication causes unrepairable DNA strand breaks, leading to increases in cell death and marked effects on replicating cells (PERSE 2021). As a standard strain, NSG shows efficacy with Cisplatin, with unsurprising efficacy observed in NR mice due to the shared genetic background of these models. Predictably, this same compound had minimal effect on B6R mice, as B6R models are known to exhibit slow tumor growth kinetics since these models form an abundance of collagen deposits which inhibit tumor growth (SARGENT *et al*. 2022). Surprisingly, Cisplatin treatment in 129R mice enhanced tumor growth, although the mechanisms responsible for this phenotype have yet to be elucidated. It is possible that 129R mice have reduced or slowed Cisplatin absorption, and thus these observations are of a hormetic effect, or some other mechanism entirely. Regardless of the mechanism, these findings underscore the importance of including genetic differences in preclinical research and emphasize the need for caution in extrapolating the efficacy of a therapy in a single genetic model to inherently heterogeneous human populations. Further, our findings suggest that the tumor microenvironment may be essential in determining the fate of a new molecule or combination of compounds, posing an important consideration for therapy selection in patient care.

Our proteomic studies have shown that the same cell line xenografted in one mouse strain results in strikingly different proteomic profiles when xenografted into another strain (HASHAM *et al*. 2024). These data allude to cancer mechanisms promoting adaptation in order to optimize growth in a given environment, or to the selection for a subclone that performs favorably in that specific microenvironment. Alternatively, one could speculate that the cancer cell line irreversibly changes to suit a certain type of microenvironment and remains as such. However, our results do not support the adaptation hypothesis since tumors grown in an inhibitive microenvironment did not adapt to improved growth when introduced to inducive microenvironment, and vice versa. It is well established that clonal selection promotes growth within a tumor, but here we show evidence that this selection occurs independently of the immune system, a finding which cannot be clearly elucidated when studying patient samples directly, since patients still have a somewhat functional immune system.

Overall, our findings in genetically distinct models support the concept that the genetic composition of the tumor microenvironment plays a critical role not only in modulating cancer growth kinetics, but also in determining a host’s capacity to sustain cancers and respond to therapy. These results highlight the importance of considering the genetic background differences of the host when selecting experimental models for studying cancer biology and evaluating therapeutic efficacy. Neglecting genetic diversity in preclinical cancer models can lead to the premature dismissal of effective therapies or the costly advancement of treatments that only work in narrow genetic contexts, thereby fueling clinical trial failures and stalling meaningful progress in therapeutic development. Strain selection should not be viewed as a trivial detail, as it can profoundly influence the outcomes and translatable potential of preclinical investigations.

## Materials and Methods

### Mice

All mice were obtained from The Jackson Laboratory (JAX). JAX has 3 three of these lines readily available, BALB/cJ *Rag1−/−* (JR003145) (BAR), C57BL/6J *Rag1−/−* (JR002216) (B6R), and NOD/ShiLtJ *Rag1−/−* (JR003729) (NR). For *Rag1* inactivation in the other strains, we used a CRISPR/Cas9 Base Editor system, as described in *DMM* Sargent et al (2022) (SARGENT *et al*. 2022) to generate 129S1/SvImJ-Rag1<em3mvw>/Mvw (129R), A/J-Rag1<em2mvw>/Mvw (A/JR), CAST/EiJ-Rag1<em6mvw>/Mvw (CASTR), and PWK/PhJ-Rag1<em5mvw>/Mvw (PWKR). NSG (JR005557) and J:NU (JR007850) control animals were also purchased from JAX. *Rag1-/-Il2rg-/-* mice were established in the wild derived strains CAST/EiJ and PWK/PhJ, developing strains deficient in B- and T-lymphocytes and NK cells. These double knockout strains are CAST/EiJ-Rag1<em6mvw> Il2rg<em2mvw>/Mvw (CRG) and PWK/PhJ-Rag1<em5mvw> Il2rg<em1mvw>/Mvw (PRG), respectively. Additionally, we used the readily available NSG and NRG (JR007799) mice, which are derived from the NOD/ShiLtJ inbred parental line available from JAX, permitting assessment of the impact of NK cells on cancer growth in mice of the same genetic background.

### Animal Use

All protocols and experiments performed were approved by JAX’s Institutional Animal Care and Use Committee (IACUC), and all regulations and accreditations are approved by the American Association for Accreditation of Laboratory Animal Care (AAALAC). The Comparative Medicine and Quality (CMQ) team advances and protects the health, welfare, and genetic quality of animals at JAX. Animal Welfare and Compliance assures that JAX routinely meets the highest standards of animal care and humane treatment; and coordinates the activities of the IACUC, which oversees all aspects of the care and use of animals at the Laboratory, including training in biomethodology and surgery. JAX Veterinary Services provide clinical veterinary care for the animals, animal care oversight, and surgically altered animals on request. All mice, *Mus musculus musculus, Mus musculus domesticus,* and *Mus musculus castneus,* were obtained and reproduced at JAX (Bar Harbor, ME, USA). Mice used were between the ages of 8 and 12 weeks of age when xenografted and housed in disposable boxes with alpha pad bedding, except J:NU which require alpha-dri bedding. The mice were kept in a pressurized individual ventilated rack in an Animal Biosafety Level 2 (ABSL2) mouse room space. All breeding units were in a non-ABSL2 mouse room location, separate from where procedures were performed. All mice had free access to 6% fat sterilized food and acidified water, and boxes were changed weekly.

### Cladogram Generation

The cladogram was made based on a previous publication and generated by R studio version 4.3.1 (PETKOV *et al*. 2004; DUMONT *et al*. 2024)

### Cell lines used

CCRF-SB (RRID:CVCL_1860), an acute lymphoblastic leukemia cell line (from American Tissue Culture Collection, ATCC Cat no. CCL-120). MEC1 (RRID:CVCL_1870), a chronic lymphocytic leukemia human cell line, with a luciferase cassette. MDA-MB-231 Luc (RRID:CVCL_JZ05), a triple negative breast cancer cell line carrying a luciferase cassette. These cell lines were selected due to several studies that have been published using these, as well as our own studies with consistent results. The addition of a luciferase cassette allows for imaging with quantitative results to determine cancer burden compared to survival curves alone.

### Isoflurane anesthesia

A calibrated Vet Equip precision vaporizer utilized only for isoflurane was used with a compressed oxygen cylinder that was regulated to reduce the high-pressure oxygen to 50 psi, per manufacturer’s instructions. A charcoal canister was used to scavenge the isoflurane from the induction chamber and breathing circuit. Before use, the canister was weighed to ensure that the isoflurane would be scavenged properly per the manufacturer’s instructions. If the weight increase over initial weight was over the collection limits, the canister had to be replaced. All tubing and connections were checked regularly to ensure isoflurane was being delivered to the induction chamber and the nose cone. The oxygen tank required a minimum of 500 psi to ensure sufficient oxygen supply for the duration of the procedures.

Once set up, a supplemental heat source was provided to the mouse for procedures exceeding 15 minutes. To avoid overheating, a thermometer was attached to the workspace and monitored for a temperature range of 80-100 °F (26.6-38 °C). Oxygen flow was set to 1 liter/min (LPM) and adjusted to desired number setting using the flow meter knob. The mouse was placed into the induction chamber, and the lid was closed. The vaporizer dial was set to 2-2.5% and the mouse’s respiratory rate was monitored to ensure a proper depth of anesthesia. A proper depth was achieved when breathing rate reduced to approximately 1 breath per second, and the breathing rate did not change when the mouse was manipulated by toe pinch or surgical manipulation.

Once the mouse was unresponsive by toe pinch, ophthalmic ointment was applied to the eyes to prevent the cornea from drying out. The mouse was then moved to the nose cone that was set up on a heating pad and placed under a sterile drape if undergoing surgery. Another toe pinch was performed to ensure the depth of anesthesia was proper before going forward with any procedure. The mouse was observed throughout the procedure, and the isoflurane level was adjusted as needed to maintain the appropriate depth of anesthesia.

### Anesthesia recovery

When the procedure was completed, mice were placed in lateral or ventral recumbency in a clean recovery cage with grain and water and placed directly on a warming pad on one side with the bedding moved to the opposite side if alpha-dri was used. The temperature of the cage was monitored with a thermometer and maintained at 80-100 °F (26.6-38 °C) at the level of the cage floor. The recovery cage was placed with only one half of the cage on the heating pad to allow the animals access to move away from the heat source. The mice were recovered in a quiet designated area and were monitored a minimum of every 15 minutes by either the surgeon or a trained technician to ensure a full recovery. The animals were returned to the animal room once all were able to walk around the cage with a normal gait and food and water intake was observed.

### Intracardiac injections

Mice were anesthetized per the isoflurane anesthesia method described above, and fur was removed using clippers from the area around the injection site on the chest. Excess fur was removed by wiping with dry paper towel. Carprofen (10 mg/kg - 0.2mL per 10 grams of mouse, CAS no. 53716-49-7, obtained from JAX Pharmacy) was injected subcutaneously by body weight per manufacturer dosing instructions. The mouse was placed in dorsal recumbency with the nose in an anesthesia nose cone. The skin around the proposed injection site was aseptically prepared with a sterile swab soaked in 70% ethanol, starting in the middle of the proposed injection site and radiated out in widening circles. This process was repeated with chlorhexidine twice. Cells suspended in up to 100 µL of sterile DPBS (Catalog No. 14190136, Gibco ordered from ThermoFisher) were loaded into a fresh sterile syringe with a 26-gauge needle, ensuring there were no air bubbles. The needle was carefully inserted into the left thorax at the second intercostal space, approximately 3 mm lateral of the sternum and directed toward midline into the left ventricle. This was confirmed by bright red blood entering the needle hub, indicating it is oxygenated and in the left ventricle. The cells were slowly injected, over the course of 20 to 40 seconds, then the needle was withdrawn. A fresh, sterile syringe and needle were required for each mouse to avoid contamination due to blood entering the needle hub.

### Xenografting via tail vein

Mice were xenografted via tail vein using a preheated brass tail vein holder. A light was placed to keep the animal warm. The tail was cleaned with ethanol and 100 µL cells were injected using a 27-gauge needle, as previously published (SARGENT *et al*. 2022)

### Xenografting MDA-MB-231 subcutaneously

Mice were xenografted with 100 µL MDA-MB-231 subcutaneously. Mice were anesthetized with isoflurane. The fur was removed and skin was sterilized with ethanol and 2% chlorhexidine prior to injection, as previously published (SARGENT *et al*. 2022).

### Tumor Removal and Passaging

Once MDA-MB-231 Luc tumors in NR mice reached a volume of 1000 mm³, all mice were euthanized. Fur around the tumor was clipped and the entire flank and surrounding area were sterilized using 70% ethanol, as described above. An incision was made through the skin directly adjacent to the tumor, and the tumor was subsequently extracted using clean forceps and a clean set of dissecting scissors. A washing media was made using 25 mL of sterile DPBS and 300 µL of pen strep (Catalog No 10378016, Gibco – ordered from ThermoFisher). The solution was made in 75 mL batches and filtered through a 0.7 micrometer syringe filter attached to a 25 mL syringe. The tumor was then placed into a 50 mL Eppendorf tube and 25 mL of washing media was added. The tube was shaken and washing media was removed immediately. The tumor was transferred to a clean 50 mL tube and 25 mL of fresh washing solution was added. This process was repeated three times total per tumor. Once all tumors from both NR and B6R were removed and cleaned, the NR tumors were pooled together into one petri dish and the B6R tumors were pooled together in another petri dish. Necrotic tissue was removed from all tumors, and the tumor masses were measured and recorded based on the strain from which they originated. B6R had the smallest tumor volumes, with the total weight of all tumors weighing 480 g. NR tumors were much larger in volume; in order to keep the same concentration of tumor to volume ratio, 480 g of NR tumor were separated for implantation into the corresponding mice (**Figure 3A**). B6R tumors were added to 3.55 mL DPBS and minced using sterile scissors. The same procedure was performed for NR tumors. The tumors were minced to the consistency that was viscous enough to fit into a 19-gauge needle, and they were then implanted via subcutaneous injection method as described previously using fresh 19-gauge needles (SARGENT *et al*. 2022).

### *In vivo* imaging of mice for bioluminescence

Mice were anesthetized and imaged as previously published (SARGENT *et al*. 2022) using the IVIS Spectrum CT. Images of bioluminescence signal was captured, and the max radiance was measured to determine the amount of cancer that engrafted into the animal (SARGENT *et al*. 2022). Image analysis was completed using Living Image version 4.8.2.

## Statistical Methods

All statistics were calculated using GraphPad Prism Version 9.4.1 (458) as indicated in the figure legends. Two-way ANOVA was used to calculate P-values for all, the data.

## Acknowledgements

We thank our scientific writer, Dr. Cara Hardy, for editing and critiquing the manuscript. We thank Drs. Steven Munger and Julie Wells for their feedback on the data. We acknowledge our summer student Ashley Kuczmera and JAX Education for contributing to this article (*Summer Student Program 2024*). We thank Dr. Michael Wiles (formerly the Technology Evaluation Department) for providing us with the CASTR, PWKR, CRG, and PRG mice. We would like to thank Jennifer Norman and Olivier Donzé for the InVivoKine samples.

We acknowledge the following JAX Scientific Research Services for their assistance in this study: The Xenografting and Live Imaging Core, Library Sciences (Jason Felty), and Scientific Instrument Services (Jeff Stone). MDA-MB-231-Luciferase cells and MEC1-Luciferase cells are kind gifts from Dr. Karolina Palucka (JAX; for references, see (SARGENT *et al*. 2022)), and Dr. Kevin Mills (formerly at JAX; (WILSON *et al*. 2019)) respectively. These cell lines have been described and validated in the references above.

## Author contributions

Conceptualization: M.G.H.; Methodology: J.K.S., M.A.W., S.R.F, B.L.D.; Formal analysis: B.L.D., M.G.H., Writing-original draft: M.A.W., M.G.H.; Writing - review & editing: J.K.S., M.A.W., S.R.F, B.L.D, M.G.H.; Supervision: M.G.H.

## Data and resource availability

All relevant data and details of resources can be found within the article and its supplementary information. Additional data is available upon request.

---

**Fig. S1.**
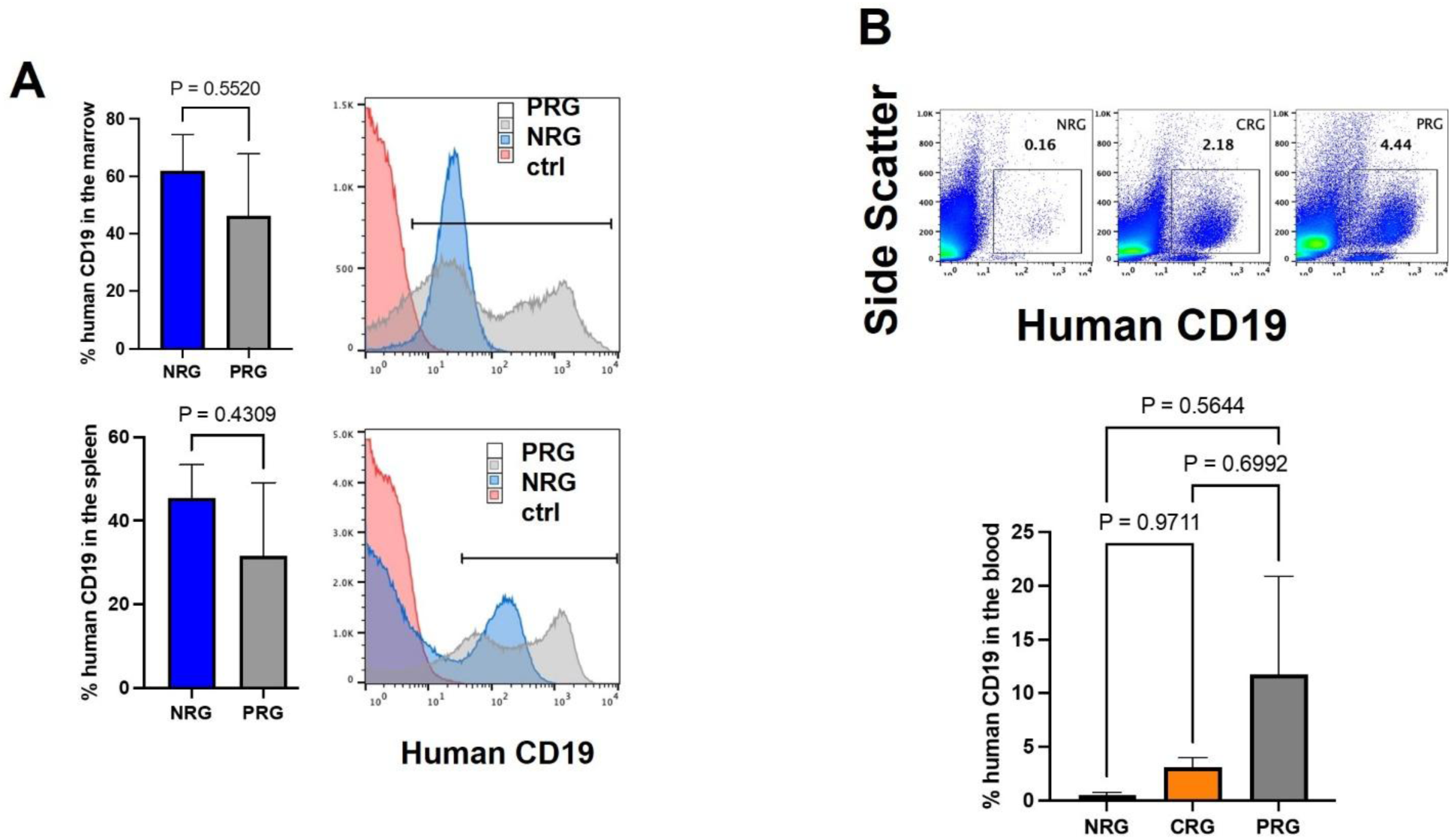
Flow cytometry results of human CD19 expression: **(A)** MEC1 in the spleen and bone marrow and **(B)** CCRF-SB cells in the blood

**Fig. S2.**
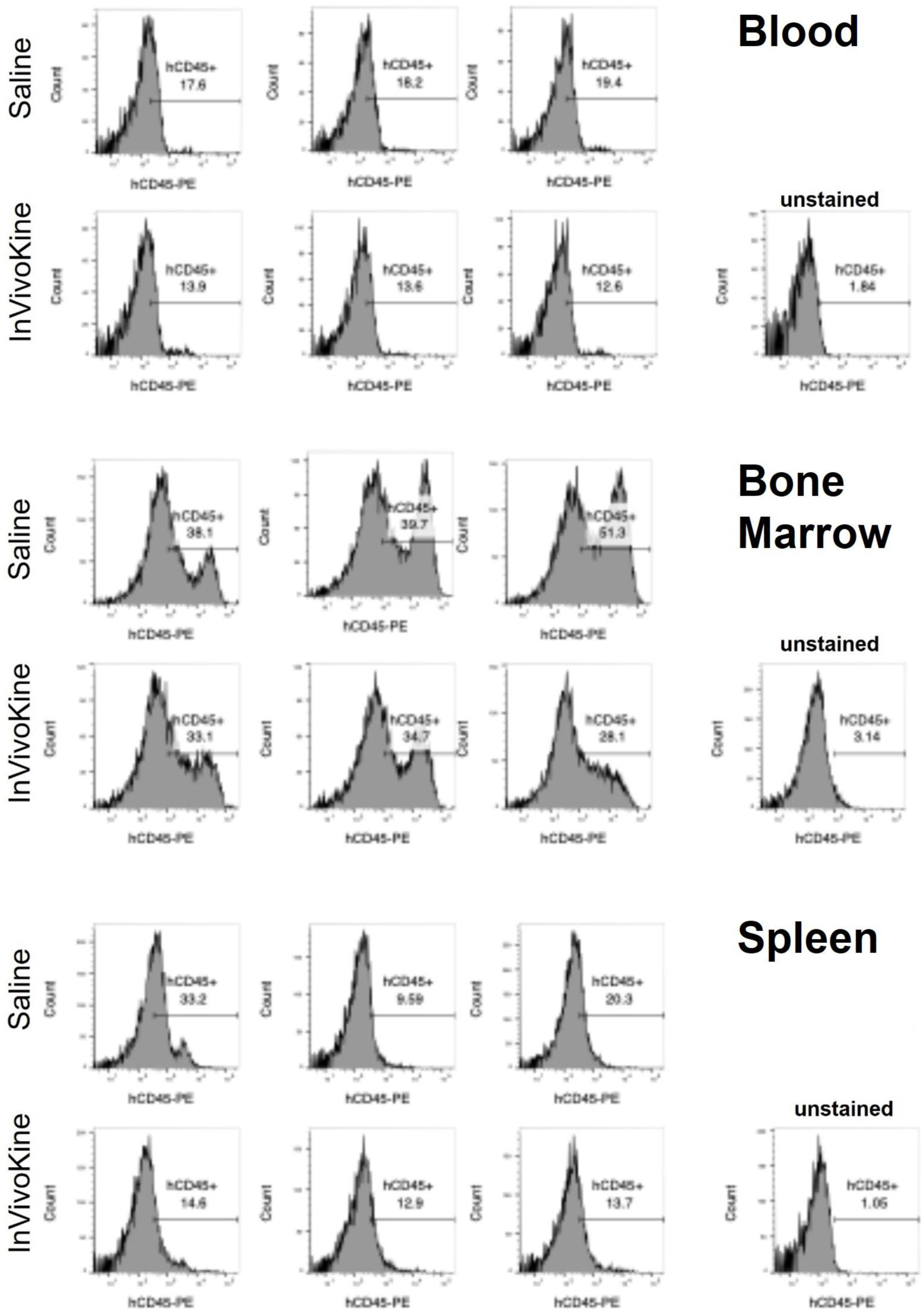
Flow cytometry results of human CD45 expression: Individual results of human CD45 expression in CCRF-SB injected mice comparing treatment with IL-1Ra InVivoKine vs saline control.

## REFERENCES

Anderson, N. M., and M. C. Simon, 2020 The tumor microenvironment. Curr Biol 30: R921–R925.

Arnold, M., M. S. Sierra, M. Laversanne, I. Soerjomataram, A. Jemal et al., 2017 Global patterns and trends in colorectal cancer incidence and mortality. Gut 66: 683–691.

Bosenberg, M., E. T. Liu, C. I. Yu and K. Palucka, 2023 Mouse models for immuno-oncology. Trends Cancer 9: 578–590.

Cao, X., E. W. Shores, J. Hu-Li, M. R. Anver, B. L. Kelsall et al., 1995 Defective lymphoid development in mice lacking expression of the common cytokine receptor gamma chain. Immunity 2: 223–238.

Cheon, D. J., and S. Orsulic, 2011 Mouse models of cancer. Annu Rev Pathol 6: 95–119.

Ciantra, Z., V. Paraskevopoulou and I. Aifantis, 2025 The rewired immune microenvironment in leukemia. Nat Immunol 26: 351–365.

Coate, L., S. Cuffe, A. Horgan, R. J. Hung, D. Christiani et al., 2010 Germline genetic variation, cancer outcome, and pharmacogenetics. J Clin Oncol 28: 4029–4037.

de Visser, K. E., and J. A. Joyce, 2023 The evolving tumor microenvironment: From cancer initiation to metastatic outgrowth. Cancer Cell 41: 374–403.

Dumont, B. L., D. M. Gatti, M. A. Ballinger, D. Lin, M. Phifer-Rixey et al., 2024 Into the Wild: A novel wild-derived inbred strain resource expands the genomic and phenotypic diversity of laboratory mouse models. PLoS Genet 20: e1011228.

Elhanani, O., R. Ben-Uri and L. Keren, 2023 Spatial profiling technologies illuminate the tumor microenvironment. Cancer Cell 41: 404–420.

Fugmann, S. D., 2001 RAG1 and RAG2 in V(D)J recombination and transposition. Immunol Res 23: 23–39.

Hanahan, D., 2026 Hallmarks of cancer-Then and now, and beyond. Cell.

Hasham, M. G., J. K. Sargent, M. A. Warner, S. R. Farley, B. R. Hoffmann et al., 2024 Methods to study xenografted human cancer in genetically diverse mice. bioRxiv.

Hoffman, R. M., 2017 Patient-Derived Mouse Models of Cancer: Patient-Derived Orthotopic Xenografts (PDOX), pp. 1 online resource (XVI, 296 pages 260 illustrations, 250 illustrations in color in Molecular and Translational Medicine,. Springer International Publishing: Imprint: Humana,, Cham.

Holland, E. C., 2004 Mouse models of human cancer. Wiley-Liss, Hoboken, N.J.

Hopken, U. E., and A. Rehm, 2019 Targeting the Tumor Microenvironment of Leukemia and Lymphoma. Trends Cancer 5: 351–364.

Jung, J., H. S. Seol and S. Chang, 2018 The Generation and Application of Patient-Derived Xenograft Model for Cancer Research. Cancer Res Treat 50: 1–10.

Olson, B., Y. Li, Y. Lin, E. T. Liu and A. Patnaik, 2018 Mouse Models for Cancer Immunotherapy Research. Cancer Discov 8: 1358–1365.

Parham, P., and C. Janeway, 2015 The immune system. Garland Science, Taylor & Francis Group, New York, NY.

Pektas, G., E. Gonul, S. Oncu, M. Becit Kizilkaya, G. Sadi et al., 2025 Chronic Lymphocytic Leukemia: Investigation of Survival and Prognostic Factors with Drug-Related Remission. Diagnostics (Basel) 15.

Perse, M., 2021 Cisplatin Mouse Models: Treatment, Toxicity and Translatability. Biomedicines 9.

Petkov, P. M., Y. Ding, M. A. Cassell, W. Zhang, G. Wagner et al., 2004 An efficient SNP system for mouse genome scanning and elucidating strain relationships. Genome Res 14: 1806–1811.

Phifer-Rixey, M., B. Harr and J. Hey, 2020 Further resolution of the house mouse (Mus musculus) phylogeny by integration over isolation-with-migration histories. BMC Evol Biol 20: 120.

Prabhakaran, P., F. Hassiotou, P. Blancafort and L. Filgueira, 2013 Cisplatin induces differentiation of breast cancer cells. Front Oncol 3: 134.

Roberts, A., F. Pardo-Manuel de Villena, W. Wang, L. McMillan and D. W. Threadgill, 2007 The polymorphism architecture of mouse genetic resources elucidated using genome-wide resequencing data: implications for QTL discovery and systems genetics. Mamm Genome 18: 473–481.

Sargent, J. K., M. A. Warner, B. E. Low, W. H. Schott, T. Hoffert et al., 2022 Genetically diverse mouse platform to xenograft cancer cells. Dis Model Mech 15.

Sung, H., J. Ferlay, R. L. Siegel, M. Laversanne, I. Soerjomataram et al., 2021 Global Cancer Statistics 2020: GLOBOCAN Estimates of Incidence and Mortality Worldwide for 36 Cancers in 185 Countries. CA Cancer J Clin 71: 209–249.

Talseth-Palmer, B. A., and R. J. Scott, 2011 Genetic variation and its role in malignancy. Int J Biomed Sci 7: 158–171.

Thakkar, S., D. Sharma, K. Kalia and R. K. Tekade, 2020 Tumor microenvironment targeted nanotherapeutics for cancer therapy and diagnosis: A review. Acta Biomater 101: 43–68.

Uthamanthil, R., P. Tinkey and E. De Stanchina, 2017 Patient derived tumor xenograft models: promise, potential and practice. Elsevier/AP, Academic Press is an imprint of Elsevier, Amsterdam.

Wilkinson, L., and T. Gathani, 2022 Understanding breast cancer as a global health concern. Br J Radiol 95: 20211033.

Wilson, J. J., K. H. Chow, N. J. Labrie, J. A. Branca, T. J. Sproule et al., 2019 Enhancing the efficacy of glycolytic blockade in cancer cells via RAD51 inhibition. Cancer Biol Ther 20: 169–182.

Wu, T., and Y. Dai, 2017 Tumor microenvironment and therapeutic response. Cancer Lett 387: 61–68.

Xiao, Y., and D. Yu, 2021 Tumor microenvironment as a therapeutic target in cancer. Pharmacol Ther 221: 107753.

